# “Theory, practice, and conservation in the age of genomics: the Galápagos giant tortoise as a case study”

**DOI:** 10.1101/187534

**Authors:** Stephen J Gaughran, Maud C Quinzin, Joshua M Miller, Ryan C Garrick, Danielle L Edwards, Michael A Russello, Nikos Poulakakis, Claudio Ciofi, Luciano B Beheregaray, Adalgisa Caccone

**Affiliations:** Department of Ecology and Evolutionary Biology, Yale University, 21 Sachem St. New Haven, Connecticut, 06520, United States of America; Department of Biology, University of Mississippi, Oxford, Mississippi, 38677, United States of America; Life and Environmental Sciences, University of California, Merced, 5200 N Lake Rd, Merced, California, 95343, United States of America; Department of Biology, University of British Columbia, Okanagan Campus, Kelowna, BC V1V 1V7, Canada; Department of Biology, School of Sciences and Engineering, University of Crete, Vasilika Vouton, Gr-71300, Heraklio, Crete, Greece; Natural History Museum of Crete, School of Sciences and Engineering, University of Crete, Knossos Av., GR-71409, Heraklio, Crete, Greece; Department of Biology, University of Florence, 50019 Sesto Fiorentino (FI), Italy; Molecular Ecology Lab, School of Biological Sciences, Flinders University, GPO Box 2100, Adelaide, SA, 5001, Australia

## Abstract

High-throughput DNA sequencing allows efficient discovery of thousands of single nucleotide polymorphisms (SNPs) in non-model species. Population genetic theory predicts that this large number of independent markers should provide detailed insights into population structure, even when only a few individuals are sampled. Still, sampling design can have a strong impact on such inferences. Here, we use simulations and empirical SNP data to investigate the impacts of sampling design on estimating genetic differentiation among populations that represent three species of Galápagos giant tortoises (*Chelonoidis* spp.). Though microsatellite and mitochondrial DNA analyses have supported the distinctiveness of these species, a recent study called into question how well these markers matched with data from genomic SNPs, thereby questioning decades of studies in non-model organisms. Using >20,000 genome-wide SNPs from 30 individuals from three Galápagos giant tortoise species, we find distinct structure that matches the relationships described by the traditional genetic markers. Furthermore, we confirm that accurate estimates of genetic differentiation in highly structured natural populations can be obtained using thousands of SNPs and 2-5 individuals, or hundreds of SNPs and 10 individuals, but only if the units of analysis are delineated in a way that is consistent with evolutionary history. We show that the lack of structure in the recent SNP-based study was likely due to unnatural grouping of individuals and erroneous genotype filtering. Our study demonstrates that genomic data enable patterns of genetic differentiation among populations to be elucidated even with few samples per population, and underscores the importance of sampling design. These results have specific implications for studies of population structure in endangered species and subsequent management decisions.

*“Modern molecular techniques provide unprecedented power to understand genetic variation in natural populations. Nevertheless, application of this information requires sound understanding of population genetics theory.”*

*- Fred Allendorf (2017, p. 420)*

## Introduction

The advent of high-throughput DNA sequencing has enabled the characterization of the genomes of model and non-model organisms alike. Genome-wide data can improve the precision and accuracy of estimates of population parameters, enhancing our understanding of present-day structure, gene flow, and local adaptation (Funk *et al.* 2012). These data have also facilitated more detailed reconstructions of historical events that impacted evolutionary trajectories within species (e.g., Emerson et al. 2010), and among closely related species (e.g., Chaves et al. 2016).

While whole genome sequencing is still beyond the budget of many research programs, methods based on reduced representation genomic libraries (e.g., double digest Restriction-site Associated DNA sequencing, ddRADseq (Peterson *et al.* 2012)) allow tens or hundreds of thousands of single nucleotide polymorphisms (SNPs) to be discovered and reliably genotyped at a much-reduced cost (Andrews *et al.* 2016). This is particularly beneficial for species of conservation concern, where limited resources and sampling constraints (i.e., few individuals are available) may be prevalent. No matter the application, though, well-designed population genetics studies aim to maximize their statistical power while minimizing costs.

Genome-wide SNP data are currently being applied to a broad spectrum of conservation objectives. These range from informing captive breeding programs (e.g., Wright et al. 2015) and improving detection of hybridization and inbreeding depression (e.g., Robinson et al. 2016; vonHoldt et al. 2016b), to delineating conservation units, assessing levels of adaptive genetic variation, and predicting viability in the face of anthropogenic impacts such as climate change (Henry & Russello 2013; Rellstab *et al.* 2015; Sork *et al.* 2016; Brauer *et al.* 2016). The appeal of genomic approaches to conservation biology is heightened by indications that a large number of independent loci can alleviate issues associated with small sample sizes per population; when using thousands of loci one can obtain reliable estimates of genetic diversity and population differentiation, so long as the true values of these parameters are sufficiently high (e.g., Li and Durbin 2011; Willing et al. 2012). Yet, as noted by Allendorf (2017), genomic datasets need to be analyzed within the context of a carefully considered sampling design. Shortcomings in sampling design can lead to erroneous conclusions (Meirmans 2015), which can have profound consequences for any population level study, but especially for those with direct management implications for threatened or endangered species.

Here, we explore the power of using thousands of SNP markers to study population structure, and the impact of sampling design and small sample sizes on detecting and describing that structure. To do this, we use genomic data from Galápagos giant tortoises (*Chelonoidis* spp.) as a case study, given a recent study has questioned the genomic distinctiveness of several species within this genus (Loire *et al.* 2013). The Galápagos Islands are home to a radiation of endemic giant tortoises that includes 11 endangered and 4 extinct species (Fig. 1). Taxonomic designations are supported by differences in morphology, geographic isolation of most species, and evidence of evolutionary divergence based on mitochondrial DNA (mtDNA) and nuclear microsatellite data ((Ciofi et al. 2002; Beheregaray et al. 2003a; Garrick et al. 2015); see Fig. S7 A and B).

**Figure 1.**
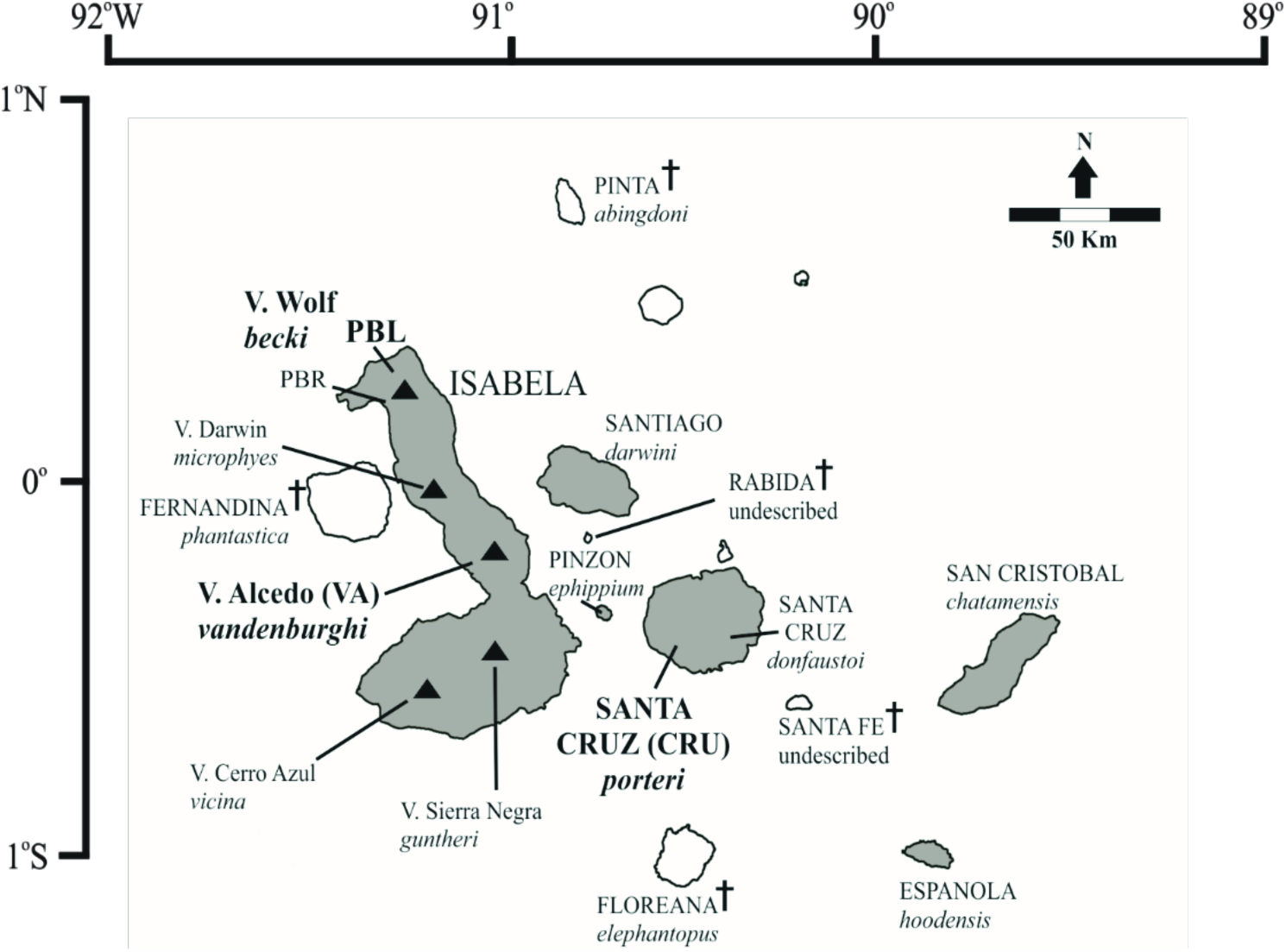
Distribution map of Galápagos giant tortoises throughout the archipelago. The islands with extant species are shown in gray, while the islands with extinct species are in white. Black triangles identify the location of the four volcanoes on Isabela Island, each with its own locally endemic tortoise species. Extinct species are identified by a cross symbol. Names of each species are in cursive with a black line pointing to the island or location within an island where they occur. The populations from the three species in this study are identified by two or three letter symbols in bold: CRU = *C. porteri*, Santa Cruz Island (La Caseta). VA = *C. vandenburghi*, Volcano Alcedo, central Isabela Island, and PBL = *C. becki*, Piedras Blancas, Volcano Wolf, northern Isabela Island.

In contrast to previous studies (see supplementary material section VIII for details; Ciofi et al. 2002; Beheregaray et al. 2003a; Beheregaray et al. 2004; Russello et al. 2005; Poulakakis et al. 2012; Garrick et al. 2015; Poulakakis et al. 2015), Loire et al. (2013) challenged the genetic distinctiveness of three Galápagos giant tortoise species. Those authors collected transcriptome-derived genotypic data from ~1000 synonymous SNPs from five captive individuals representing three species (*C. becki*, *C. porteri* and *C. vandenburghi*). They did not detect significant differentiation, as measured by F_ST_, when comparing two groups (one group of three *C. becki* individuals, and the second group consisting of two individuals, one *C. porteri* and one *C. vandenburghi*). These two groups were constructed on what the authors identified as natural partitions, based on the observation that their samples fall into two different mtDNA clades (Fig. S6a; Poulakakis et al. 2012). Furthermore, Loire et al. (2013) did not detect homozygosity excess, as measured by F_IT_, for which positive values would indicate population structure. Given that previous population genetic studies have largely relied upon data from mtDNA and microsatellites, such a discrepancy between these traditional markers and genomic SNPs could have wide-ranging implications, beyond the case of Galápagos giant tortoises, and therefore warrants further investigation.

In this study, we investigate the agreement of population structure analyses based on genome-wide SNPs compared to those based on mtDNA sequences and microsatellite genotypes. To do this we generated a dataset of tens of thousands of genome-wide SNPs from 30 individuals representing the same three species (*C. becki*, *C. porteri*, and *C. vandenburghi*) considered by Loire et al. (2013). Since these species form a recently diverged species complex, we treat each species as a population to compare against the null hypothesis that all Galápagos giant tortoises belong to a single species with one panmictic population. First, we address whether or not there is significant genomic differentiation among these three Galápagos giant tortoise species using newly generated SNPs. Then, we subsample our data to explore the effects of using only a few individuals per population and of pooling individuals from different populations on estimating genetic differentiation. From these subsampling simulations, we predict the range of *F*_ST_ estimates expected when using the sampling scheme of Loire et al. (2013). Finally, we reanalyze the raw RNA-seq data from Loire et al. (2013) to test our prediction.

## Materials and Methods

### Sampling and sequencing

Samples were obtained during previously conducted collection expeditions (Caccone et al. 1999; Caccone et al. 2002; Ciofi et al. 2002; Beheregaray et al. 2003a; Beheregaray et al. 2003b; Beheregaray et al. 2004; Russello et al. 2005; Ciofi et al. 2006; Russello et al. 2007; Poulakakis et al. 2008; Garrick et al. 2012; Poulakakis et al. 2012; Edwards et al. 2013; Edwards et al. 2014; Garrick et al. 2014). Approximately ten samples per population for each extant species (*n*=121 individuals in total) were selected for sequencing as part of a larger project on the phylogeography of Galápagos giant tortoises. These individuals were chosen as they displayed concordant and unambiguous genetic assignments between mitochondrial (control region, mt*CR*) and microsatellite (12 loci) ancestry based on a published database of 123 mitochondrial haplotypes (Poulakakis *et al.* 2012) and 305 genotyped individuals (Edwards *et al.* 2013) that include all the extant and extinct populations and species.

DNA was extracted from blood samples using a DNeasy Blood and Tissue kit (Qiagen) according to the manufacturer’s instructions. We then prepared ddRAD libraries following Peterson et al. (2012). For each sample, 500 ng of genomic DNA was digested with the restriction enzymes *MluCI* and *NlaIII* (New England BioLabs), and ligated with Illumina-specific adaptors representing up to 18 unique barcodes and 2 index codes. Ligated fragments of samples were pooled into 13 libraries and size-selected to be ~310 bp (range 279 – 341bp) with a BluePippin (Sage Science). Size-selected libraries included 12 to 24 individuals and were paired-end sequenced on 13 lanes of an Illumina HiSeq 2000 at the Yale Center for Genome Analysis.

### SNP calling

We used forward and reverse reads to generate a *de novo* assembly using the pyrad v.3.0.3 pipeline (Eaton 2014). Reads were de-multiplexed and assigned to each individual based on barcodes allowing for one mismatch. We replaced base calls of Q < 20 with an ambiguous base (N) and discarded sequences containing more than four ambiguities. We used 85% clustering similarity as a threshold to align the reads into loci. We set additional filtering parameters to allow for a maximum number of SNPs to be called: retaining clusters with a minimum depth of sequence coverage (Mindepth) > 5 and a locus coverage (MinCov) > 10, a maximum proportion of individuals with shared heterozygote sites of 20% (MaxSH = p.20), and a maximum number of SNP per locus of 15 (maxSNP = 15). For subsequent analyses, we filtered this dataset using vcftools (Danecek *et al.* 2011) to generate a set of polymorphic loci (23,057 SNPs) with no missing data common to all three Galápagos giant tortoises populations of interest, abbreviated PBL, CRU, and VA and corresponding to the species *C. becki*, *C. porteri*, and *C. vandenburghi*, respectively (*n* = 10 individuals each).

### Analytical methods

*F*-statistics (*F*_IT_, *F*_IS_, global *F*_ST_, and pairwise *F*_ST_) were calculated using the diveRsity package in R (Keenan *et al.* 2013), which uses a weighted Weir and Cockerham (1984) estimator. The same package was used to assess the statistical significance of these estimates by bootstrapping across loci. Through this method we established 95% confidence intervals for each estimate, accepting as significant those that did not include 0. Pairwise *F*_ST_ calculated from thousands of subsamples of the data (described below) were carried out in vcftools (Danecek *et al.* 2011) to streamline computation. We also used vcftools to calculate the number of loci out of Hardy-Weinberg equilibrium for each population and for pooled populations.

Since *F*_ST_ estimates rely on *a priori* assignment of individuals to groups that are typically based on geographic location, we used two methods that do not have this assumption to assess patterns of differentiation among our samples. To do this, we first carried out Principal Component Analysis (PCA) on all 30 individuals, using the PLINK software (Chang *et al.* 2015). Principal components 1 and 2 were plotted against each other in R. To complement the multivariate analyses, we performed a Bayesian clustering analysis, implemented in the program STRUCTURE version 2.3.4 (Pritchard *et al.* 2000; Falush *et al.* 2003), also including all 30 individuals. STRUCTURE assumes a model with K unknown clusters representing genetic populations in Hardy-Weinberg equilibrium, and then assigns individuals to each cluster based on allele frequencies. We ran 20 repetitions of STRUCTURE for K=1-5, with a burn-in of 10,000 iterations and MCMC length of 50,000 iterations. These runs used the admixture model, correlated allele frequencies among populations, and did not assume prior population information. All other parameters were left at default values. Results were post-processed and visualized using CLUMPAK (Kopelman *et al.* 2015). We used mean log likelihood values (Pritchard *et al.* 2000) and the ΔK statistic (Evanno *et al.* 2005) to infer the best K (supplementary Figure S5). Both analyses considered the 23,057 SNPs common to all individuals.

To further assess the power of our SNPs to detect population structure, we randomly subsampled individuals from each of the species and calculated pairwise *F*_ST_ for each species using these subsamples. We tested this for per-species sample sizes of *n*=2, n=3, and n=5. This process was repeated 1,000 times for each sample size. We also carried out a similar analysis maintaining all 10 individuals per population but randomly subsampling SNPs from our dataset. For these analyses we used the following number of SNPs: 25, 50, 100, 200, 500, 1000, 5000, and 10,000. This was repeated 1,000 times for each sample size. Finally, we used a subsampling scenario that directly mimicked the one in Loire *et al.* (2013) to further evaluate the impact of limited sample sizes and pooling of samples from distinct species on *F*_ST_ estimates. As was done in Loire *et al.* (2013), we compared a set of three individuals from *C. becki* to a grouping that included one *C. porteri* plus one *C. vandenburghi* individual. To account for sample variation, we repeated this grouping process 1,000 times (described in full in supplementary material section IV).

## Results

### Tortoise samples and ddRAD-seq dataset

Our sequencing generated a total of 3,094,399,092 retained reads (approximately 15 to 58 million reads per individual) after de-multiplexing and filtering reads for quality and ambiguous barcodes and ddRAD-tags. *de novo* assembly of the data resulted in 48,004,056 ddRAD-tags (approximately 320,000-465,000 per individual). From these, we called SNPs and obtained 973,321 variable sites. We then narrowed those loci down to only loci with genotypes called in every individual in our three species data set, for a total of 23,057 SNPs. For the three species of interest the number of loci retained within populations and between populations pairs are presented in Table 1. The average coverage per locus per individual was 12X (minimum 9; maximum 15).

**Table 1.** Number of polymorphic loci present in all individuals (*n*=10 per species) used for analyses of each population (diagonal) and population pair (below diagonal).

|  | PBL<br>(*C. becki*) | CRU<br>(*C. porteri*) | VA<br>(*C. vandenburghi*) |
| --- | --- | --- | --- |
| PBL<br>(*C. becki*) | 9,580 |  |  |
| CRU<br>(*C. porteri*) | 19,654 | 11,703 |  |
| VA<br>(*C. vandenburghi*) | 13,520 | 16,432 | 5,732 |

### *F*-statistics using ddRAD-seq data

Calculation of *F*-statistics revealed values consistent with highly-structured populations (*F*_IT_ = 0.257, 95% CI: 0.251 – 0.262; *F*_IS_ = 0.079, 95% CI: 0.073 – 0.084; and global *F*_ST_ = 0.193, 95% CI: 0.189 – 0.198). Using the SNPs in common to each population pair (Table 1), we found pairwise *F*_ST_ values of 0.169 (95% CI: 0.164 – 0.174) between PBL and CRU, 0.181 (95% CI: 0.175 – 0.187) between PBL and VA, and 0.233 (95% CI: 0.226 – 0.240) between CRU and VA (Table 2). These estimates were similar to, though higher than, *F*_ST_ estimates using 12 nuclear microsatellite markers (Garrick *et al.* 2015) for these species comparisons (Table 2 and supplementary materials Table S5).

**Table 2.** Pairwise F_ST_ values between given species pairs. Above the diagonal, values calculated using our dataset of SNPs with no missing data and common to the population pair, along with 95% confidence intervals. Below the diagonal, values calculated using 12 microsatellite loci from Garrick et al. (2015) (see supplementary material section VIII). Data were obtained using 10 samples for each population (PBL, VA, CRU) for the three species.

|  | PBL<br>(*C. becki*) | CRU<br>(*C. porteri*) | VA<br>(*C. vandenburghi*) |
| --- | --- | --- | --- |
| PBL<br>(*C. becki*) | xx | 0.169<br>(0.164 – 0.174) | 0.181<br>(0.175 – 0.187) |
| CRU<br>(*C. porteri*) | 0.137 | xx | 0.233<br>(0.226 – 0.240) |
| VA<br>(*C. vandenburghi*) | 0.163 | 0.202 | xx |

### PCA and STRUCTURE

The first two principal components of the PCA showed clear differentiation among individuals from the three species. PC1 accounted for approximately 12.0% of the variation among individuals and PC2 accounted for approximately 9.3% of the variation among individuals (Figure 2). Similarly, both mean log likelihood values (Pritchard *et al.* 2000) and the ΔK statistic (Evanno *et al.* 2005) supported the existence of three distinct genetic units in the STRUCTURE analysis (supplementary Figure S5). These groups correspond to the *a priori* geographic groupings used in *F*_ST_ estimates and to the three named species. Our separate analysis of loci out of Hardy-Weinberg equilibrium (HWE), the basis for the STRUCTURE algorithm, supported these findings as well. When each species was considered separately, out of 23,057 loci PBL showed 214 out of HWE, CRU showed 124 out of HWE, and VA showed 71 out of HWE. When the CRU and VA samples were pooled, the number of loci out of HWE rose to 1326. When all three species were pooled and treated as one population, 2422 loci were found to be out of HWE.

**Figure 2.**
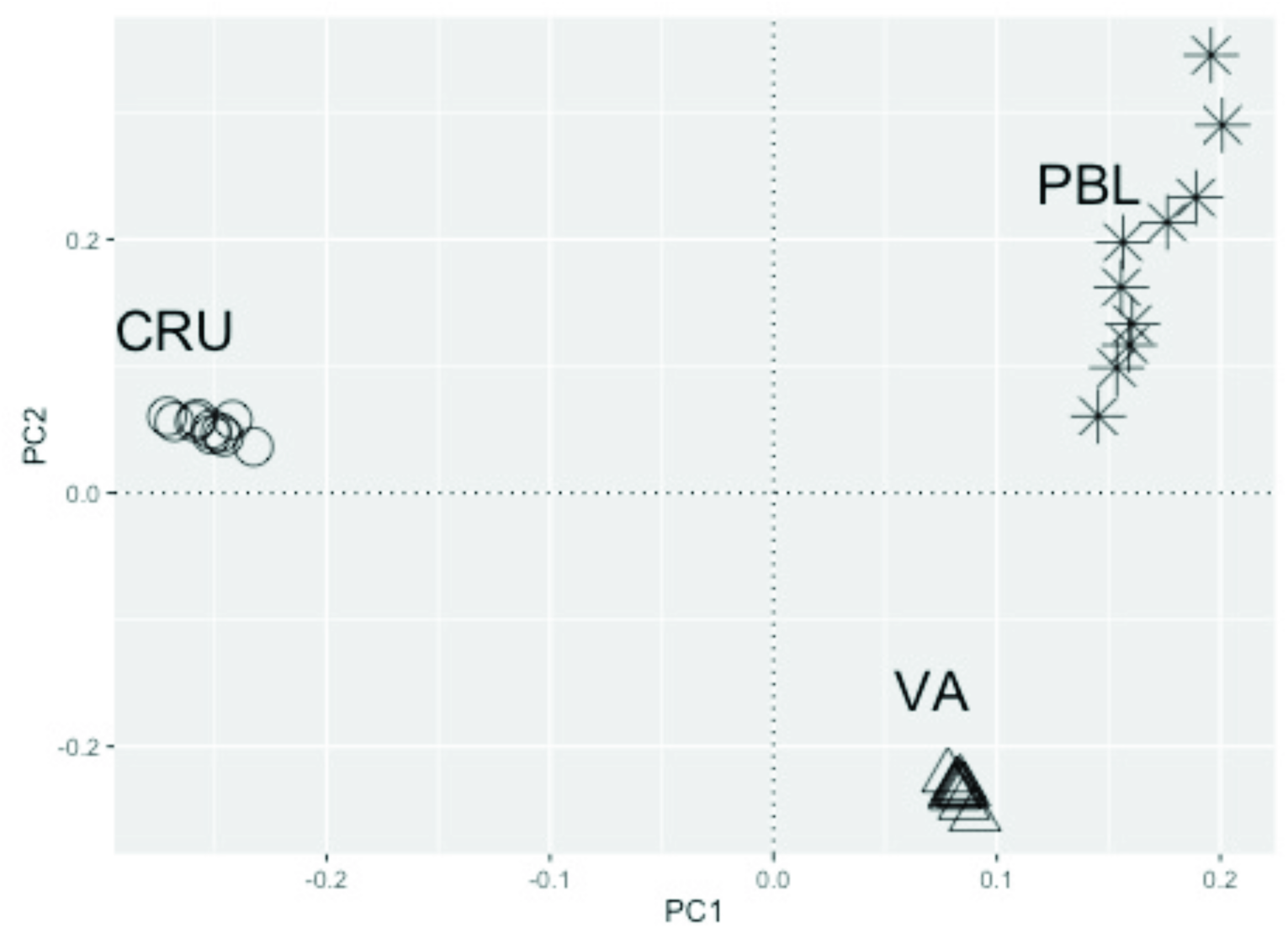
Principal component 1 (PC1) plotted against principal component 2 (PC2) for 30 individuals from 3 populations, resulting from PCA analysis on 23,057 SNPs. Stars, open circles, and open triangles identify individuals from the PBL (*C. becki*), CRU (*C. porteri*), and VA (*C. vandenburghi*) populations, respectively. The analysis was carried out using PLINK (Chang *et al.* 2015).

### Sample size, number of loci, and the effect of individual samples

In all population comparisons for the three sample sizes (n = 2, 3, or 5), the majority of estimates were within 0.03 of the *F*_ST_ value calculated using the complete dataset of 10 samples per population (Figure 3). In every case, when the sample size was two *F*_ST_ tended to be underestimated, though with a long tail of overestimated outliers. In all comparisons with sample sizes of three or five this skew disappeared: we found that 95% of the estimates were within 0.05 of the estimate using 10 samples (supplementary Tables S1).

**Figure 3.**
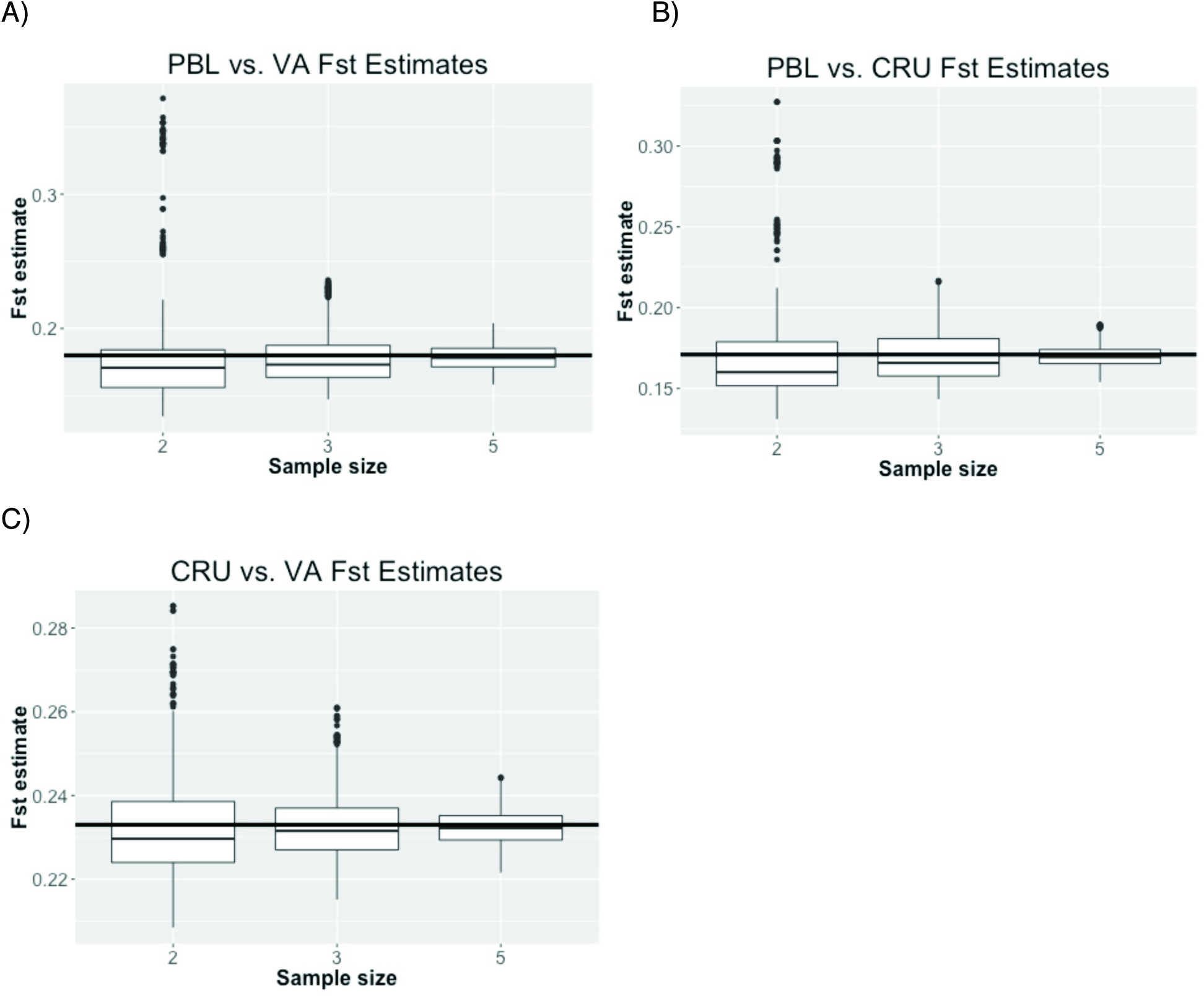
Boxplots of pairwise *F*_ST_ estimates using 1,000 randomly drawn subsamples of individuals for each sample size (*n*=2, 3, or 5) from each population. PBL, CRU and VA correspond to population samples from *C. becki*, *C. porteri* and *C. vandenburghi*, respectively. The horizontal black line in each boxplot marks the *F*_ST_ value calculated using all ten individuals from each population in the pairwise comparison (see supplementary table S1). Lower hinge corresponds to first quartile (25^th^ percentile); upper hinge corresponds to third quartile (75^th^ percentile). Whiskers indicate points within 1.5 times the interquartile range (IQR), with outliers indicated as points beyond that range.

Our *F*_ST_ estimates from subsampled SNPs ranging from 25 to 10,000 SNPs appeared to have the statistical power to detect population structure between these population pairs when 10 individuals were used, with 95% of all estimates above 0 (Table S2 A - C). However, as expected, with many fewer SNPs the range of 95% of the estimates was very wide (see supplementary material section III). For example, when only 100 SNPs were used to compare PBL and CRU, 95% of the *F*_ST_ estimates were between 0.1 and 0.255, while using 1000 SNPs gave 95% of the *F*_ST_ estimates between 0.146 and 0.194 for the same comparison (Table S2A).

### Effect of pooling samples

To test how pooling samples affected F-statistic estimates, we used the Loire *et al.* (2013) sampling design, pooling one individual from *C. porteri* and one individual from *C. vandenburghi* into one population and comparing this to three individuals from *C. becki*. When the set of common SNPs (*n*=23,057) were included in the analysis, the F_ST_ estimates between 1000 pairs of these groups ranged from 0.045 to 0.136 (95%: 0.052–0.127, mean: 0.075). When only 1,000 SNP loci were used, as in Loire *et al.* (2013), the *F*_ST_ estimates ranged from 0.006 to 0.157 (95%: 0.031–0.134, mean: 0.076) (see supplementary Figures S4A and S4B). This confirms that pooling samples from two populations, each representing different species, results in a strongly depressed *F*_ST_ estimate. However, these simulations highlight that the occurrence of genetic differentiation (i.e., positive *F*_ST_ values) should still be detectable even with this grouping scheme.

### Re-analysis of Loire et al. transcriptome data

Given that our analyses of ddRAD-seq data showed clear genetic structure among the populations from the three species, and our subsampling simulations (Figure S4) predicted that positive *F*_ST_ values should still be detectable using the grouping scheme adopted by Loire et al. (2013), we reanalyzed the original RNA-seq data generated for that publication to further assess the source of the discrepancy. We downloaded the publically-available RNA sequencing data generated by Loire et al. (2013) from the NCBI’s Sequence Read Archive and re-called SNPs after aligning these reads to a draft genome assembly of a closely related species of Galápagos giant tortoise, *C. abingdonii* (unpublished data; see methods in supplementary materials section VII). With these transcriptome-derived SNP data, we estimated an *F*_ST_ of 0.054 (95% CI: 0.049 – 0.058) when comparing the three *C. becki* samples (PBL) to the combined two *C. porteri* and *C. vandenburghi* samples (CRU and VA). Notably, this *F*_ST_ value falls within our predicted range of *F*_ST_ estimates generated by subsampling the ddRAD-seq data. Our *F*_IT_ estimate for this data set was −0.121 (95% CI: −0.129 – −0.113), with F_IS_ estimated to be −0.185 (95% CI: −0.192 – 0.177).

Plotting the first two principal components of a PCA of these five samples showed clear clustering of the conspecific samples from *C. becki*, while the single samples from *C. vandenburghi* (VA) and *C. porteri* (CRU) are distinct from each other and from the *C. becki* samples (supplementary Figure S6).

## Discussion

### Strong evidence of population structure

Using genome-wide SNP data we found evidence for significant differentiation among the three species considered (*C. becki*, *C. porteri*, and *C. vandenburghi*), consistent with the findings of decades of research in this system (Ciofi et al. 2002; Beheregaray et al. 2003a; Beheregaray et al. 2003b; Beheregaray et al. 2004; Russello et al. 2005; Russello et al. 2007; Poulakakis et al. 2008; Poulakakis et al. 2012; Garrick et al. 2015; Poulakakis et al. 2015). Our estimate of *F*_IT_ (0.257), which was a focal metric used in the previous study (Loire *et al.* 2013), was positive and significantly different from zero. Positive values of *F*_IT_ indicate an excess of homozygous loci in the sample set. This could suggest the existence of population structure in the total sample set. This possibility is reinforced by the finding of very high and significantly different from zero *F*_ST_ estimates for the same comparisons (between 0.17 and 0.24; Table 2, supplementary Figure S1). Interpreting significantly positive *F*_IS_ values, such as the one calculated from our ddRAD-seq data set, can be difficult (Allendorf and Luikart 2007). This could be due to substructure within one or more populations, sampling stochasticity, and/or recent demographic changes in relatively small populations. It could also be that such small populations are not necessarily expected to be in Hardy-Weinberg equilibrium due to the increased influence of genetic drift (Allendorf & Luikart 2009).

To assess whether there was additional genetic structure outside of our *a priori* assignment of individuals based on their geographic location, we also analyzed the 30 samples in our ddRAD-seq data set using two methods without prior assignment of each sample to a group. Both principal component (Figure 2) and Bayesian clustering analyses (supplementary Figure S5) clearly discerned three genetically distinct clusters that corresponded to the samples from the three species tested in our pairwise *F*_ST_ estimates. This echoed our per-locus analysis of Hardy-Weinberg equilibrium (HWE), which showed that treating all 30 individuals from the three named species as a single population dramatically increased the number of loci out of HWE.

Results of our analyses of population structure using tens of thousands of genome-wide SNPs are concordant with earlier studies using mtDNA haplotypes and microsatellite genotypes (Ciofi et al. 2002; Beheregaray et al. 2003a; Beheregaray et al. 2003b; Beheregaray et al. 2004; Russello et al. 2005; Russello et al. 2007; Poulakakis et al. 2008; Poulakakis et al. 2012; Garrick et al. 2015; Poulakakis et al. 2015). These findings definitively resolve concerns raised by Loire et al. (2013) regarding whether these traditional markers were accurately reflecting the genetic distinctiveness of Galápagos giant tortoise species. Importantly, our results not only revealed the same genetic clustering as earlier studies, but also showed the same patterns of genetic distance. As in the microsatellite studies, we found slightly greater genetic differentiation between *C. becki* and *C. vandenburghi* (PBL and VA: *F*_ST_ = 0.181) than between *C. becki* and *C. porteri* (PBL and CRU: *F*_ST_ = 0.169), and the greatest differentiation between *C. porteri* and *C. vandenburghi* (CRU and VA: *F*_ST_ = 0.233) (Table 2). While qualitatively the same, our *F*_ST_ estimates are notably higher than those calculated using microsatellites (Table 2), a finding predicted by the mathematics of using biallelic vs. multiallelic loci (Putman & Carbone 2014), which has also been found in other systems (e.g., Payseur and Jing 2009).

### Impact of sample size and number of loci on detecting population structure

Population genetic theory (Nei 1978), simulations (Willing *et al.* 2012), and empirical work (Reich *et al.* 2009) support the idea that a data set of thousands of loci should have the power to detect population structure with high precision, even when only a few individuals per population are analyzed. We tested this idea with our Galápagos giant tortoise ddRAD-seq SNP data by estimating *F*_ST_ from subsamples of two, three, and five individuals from each population and comparing them to the same estimates obtained from 10 individuals per population. All tested sample sizes were able to detect significant *F*_ST_ values, though using three or five samples yielded more precise estimates than using only two (Figure 3; supplementary tables S1). These analyses are consistent with the idea that accurate F_ST_ values can be estimated using as few as two or three samples per population if thousands of SNPs are analyzed. Likewise, we found that for highly differentiated populations such as those studied here, hundreds of SNPs were sufficient to accurately describe population structure when ten individuals per population were used. This empirical evidence should be helpful in the design of future conservation genetics studies that aim to describe population structure, in which case additional samples may lead to diminishing returns for improving statistical power. This will be especially useful for endangered or elusive species for which sampling may present a severe limitation.

### Sampling design matters

Our genome-wide SNP data detected high and significant differentiation among these three species, even when only two or three individuals from each were used in the analysis (Figure 3). While these results were strongly supported, they failed to explain the discrepancy described by Loire et al. (2013), who used over 1,000 synonymous SNPs from transcriptome sequencing data and found no differentiation between the same three species. Their sample size of five captive individuals does not by itself account for the discrepancy between the two studies, because, as we show above (supplementary Figure S4), using thousands of SNPs should give sufficient power to detect population structure in Galápagos giant tortoises, even when sample size is that small.

Instead, sampling design, and specifically grouping of individuals into inappropriate population units, rather than sample size likely biased the statistical power of Loire et al.’s (2013) study. Their sampling scheme divided the five individuals into two groups, which did not reflect the population divergence of the three species. Specifically, this mixed group included two individuals, each from different species (CRU, *C. porteri* from Santa Cruz Island; VA, *C. vandenburghi* from central Isabela Island), and another group of three individuals from the other species (PBL, *C. becki* from northern Isabela Island). The justification for this grouping was based on the closer phylogenetic relationship of mtDNA haplotypes from *C. porteri* and *C. vandenburghi* (Caccone et al. 1999; Russello et al. 2007) compared to haplotypes found in the PBL *C. becki* population. This choice is problematic for several reasons (detailed in the supplementary material section VIII). Most importantly, *F*-statistics are a reflection of population differentiation, not of phylogenetic relatedness. Treating the individuals from *C. porteri* and *C. vandenburghi* as belonging to the same population biased the *F*-statistics estimates by leading to an increase in within-group variation, and therefore depressed *F*_ST_ values. This within-group structure, which distorts *F*-statistics, is known as Wahlund effect (Wahlund 1928).

The problem outlined above is clear in our pairwise analysis using >20,000 SNPs, which shows that while the *C. becki* population sample is about equally differentiated from the *C. porteri* and *C. vandenburghi* ones, the ones from *C. porteri* and *C. vandenburghi* are more differentiated from each other than from the *C. becki* population sample (Table 2). To empirically test for the Wahlund effect under this sampling scheme, we simulated a scenario in which three samples from *C. becki* were compared to a population consisting of one *C. porteri* and one *C. vandenburghi* sample. Repeating this sampling scenario 1,000 times, we found significantly depressed mean F_ST_ estimates, as low as 0.075, with 95% of comparisons ranging from 0.052 to 0.127 (supplementary Figure S4A). Even more strikingly, when we limited the analysis to a similar number of markers as Loire et al. (2013) and used 1,000 randomly drawn SNPs, the range of 95% of the estimates increased to 0.031 to 0.134.

### RNA-seq data supports population structure

While our subsampling simulations showed a clear Wahlund effect when samples from two different species (*C. porteri* and *C. vandenburghi*) were combined into one grouping, these *F*_ST_ estimates were still positive (mean *F*_ST_ = 0.075). We therefore would have expected Loire et al. (2013) to find a similar estimate in their analysis of RNA-seq data, but they reported no significantly positive *F*_ST_ value. To investigate this discrepancy, we re-analyzed their raw sequencing data by aligning it to a Galápagos giant tortoise reference genome. Using the SNPs from this reanalysis, we estimated an *F*_ST_ of 0.054, which is similar to our expected *F*_ST_ under their sampling design (supplementary Figure S4). Our estimates of *F*_*IS*_ and *F*_*IT*_ for the RNA-seq data set were negative, a surprising result that may be related to the sampling design, the specific individuals included in that study, or the deviations from Hardy-Weinberg equilibrium that can occur in small populations (Kimura & Crow 1963). This last point is due to the assumption of large numbers in Hardy-Weinberg equilibrium, which is violated in small populations (Allendorf & Luikart 2009).

Convincingly, a PCA of Loire et al.’s (2013) SNP data revealed a tight cluster of the three PBL samples, whereas the CRU and VA samples were distinct both from the PBL cluster and from each other (Figure S6). This pattern of principal components mirrors the one that we found with our 30 sample dataset for the same populations (Figure 2). These results, which match our expectations based on subsampling simulations (supplementary figure S4), suggest that the lack of significantly positive *F*_ST_ values found by Loire et al. (2013) is due not just to small sample size and inappropriate grouping of samples, but also the genotype filters employed in their initial analysis. The original Loire et al. (2013) methods describe a genotype filter that assigns posterior probability to genotypes based on Hardy-Weinberg equilibrium. We suspect that this may not be a reliable method when genotyping a pool of individuals from different species, since these samples will not meet the assumption of Hardy-Weinberg equilibrium. Our SNP calls of their data may have also been improved by mapping the RNA sequence reads to a draft Galápagos giant tortoise reference genome, as suggested by others (Shafer *et al.* 2016). However, our ddRAD-seq SNP data were called without mapping to a reference, so this methodological difference cannot completely explain the loss of signal.

### Conclusions

Reduced-representation sequencing offers practical ways to take advantage of the power of population genomics, even when samples and funds are limited (Narum *et al.* 2013). Yet, thoughtful study design remains an essential component. Our analyses clearly showed that tortoises representing each of three named species exhibit high genetic differentiation at the genomic level, as demonstrated through high and significant *F*_ST_, and positive *F*_IT_ estimates, as well as through principal component and Bayesian clustering analyses. Using thousands of SNPs gives high statistical power to detect population structure even when sample sizes of individuals are as few as two or three individuals. However, the heterogeneity of samples within a population can confound calculations using small sample sizes in unpredictable ways. Reduced sample size also limits the diversity of analyses that can be performed, especially limiting those that do not rely on a priori population designation, such as principal component analysis and Bayesian clustering algorithms. Ultimately, we found that both our ddRAD-seq data and a reanalysis of RNA-seq data generated by Loire *et al.* (2013) were consistent with the findings of earlier microsatellite and mtDNA studies. We therefore expect genome-wide SNPs to support the conclusions of population genetic studies of Galápagos giant tortoises beyond the three species considered here.

Distinguishing populations and evolutionary lineages, such as the giant tortoise species analyzed here, is a vital role for population genetic analyses to play in conservation (Funk *et al.* 2012). Results from such analyses can assist in protected area designation (Larson *et al.* 2014), inform appropriate legal protections (vonHoldt et al. 2016a), and guide captive breeding strategies (de Cara *et al.* 2011; Lew *et al.* 2015). We show that, as long as population genetics theory is carefully taken into account, the use of genome-wide data enabled by high-throughput sequencing can be a powerful tool in these conservation efforts, even when sample sizes are limited.

## Data Accessibility

Raw data from Illumina sequencing will be deposited to the NCBI Short Read Archive (SRA) for all individuals included in this study. The vcf file used in the analyses will be deposited on Dryad. Microsatellite genotypes and mitochondrial DNA sequences used in the supplementary material are available upon request.

